# Metabolic signatures of birth weight in 18 288 adolescents and adults

**DOI:** 10.1101/049247

**Authors:** Peter Würtz, Qin Wang, Marjo Niironen, Tuulia Tynkkynen, Mika Tiainen, Fotios Drenos, Antti J Kangas, Pasi Soininen, Michael R Skilton, Kauko Heikkilä, Anneli Pouta, Mika Kähönen, Terho Lehtimäki, Richard J Rose, Eero Kajantie, Markus Perola, Jaakko Kaprio, Johan G Eriksson, Olli T Raitakari, Debbie A Lawlor, George Davey Smith, Marjo-Riitta Järvelin, Mika Ala-Korpela, Kirsi Auro

## Abstract

**Background:** Lower birth weight is associated with increased susceptibility to cardiometabolic diseases in adulthood, but the underlying molecular pathways are incompletely understood. We examined associations of birth weight with a comprehensive metabolic profile measured in adolescents and adults.

**Methods:** High-throughput nuclear magnetic resonance metabolomics and biochemical assays were used to quantify 87 circulating metabolic measures in seven cohorts from Finland and the United Kingdom comprising altogether 18 288 individuals (mean age 26 years, range 15–75). Metabolic associations with birth weight were assessed by linear regression models adjusted for sex, gestational age, and age at blood sampling. The metabolic associations with birth weight were compared to the corresponding associations with adult body mass index (BMI).

**Results:** Lower birth weight was adversely associated with cardiometabolic biomarkers, including lipoprotein subclasses, fatty acids, amino acids, and markers of inflammation and impaired liver function (P<0.0015 for 46 measures). Associations were consistent across cohorts with different ages at metabolic profiling, but the magnitudes were weak. The pattern of metabolic deviations associated with lower birth weight resembled the metabolic signature of higher adult BMI (*R*^2^=0.77). The resemblance indicated that 1-kg lower birth weight is associated with similar metabolic aberrations as caused by 0.92-units higher BMI in adulthood.

**Conclusion:** Lower birth weight is associated with adverse biomarker aberrations across multiple metabolic pathways. Coherent metabolic signatures between lower birth weight and higher adult adiposity suggest potentially shared underlying molecular mechanisms. However, the magnitudes of metabolic associations with birth weight are modest in comparison to the effects of adiposity, implying that birth weight is only a weak indicator of metabolic risk in adulthood.

**KEY POINTS:**

- Lower birth weight is adversely associated with a wide range of established and emerging circulating cardiometabolic biomarkers in adulthood, including lipoprotein subclasses and their lipids, fatty acid balance, amino acids, and markers of inflammation and liver function.
- The metabolic associations are consistent across a wide age span from adolescence to retirement age, and similar for men and women.
- The magnitudes of metabolic aberrations are weak for the variation in birth weight observed in general population cohorts. Although the metabolic associations with birth weight are statistically significant, they are likely to be of minor public health relevance.
- The overall metabolic association pattern with lower birth weight closely resembles the metabolic signature of higher adult adiposity, suggesting that shared underlying metabolic pathways may be involved.
- 1-kg lower birth weight (≈2 SD) is associated with similar adverse metabolic effects as caused by 0.92 higher BMI (≈0.25 SD) in adulthood. These findings indicate that fetal growth, as assessed by birth weight, only has minor effects on the adult metabolic risk profile in general population settings.

## INTRODUCTION

Low birth weight is a marker of impaired fetal growth and has been associated with increased rates of ischemic heart disease, stroke, and type 2 diabetes.^1-4^ The developmental-origins hypothesis proposes that fetal adaptive responses to undernutrition, caused by maternal nutritional or placental factors, is an underpinning factor between lower birth weight and increased susceptibility to cardiometabolic diseases.^5,6^ Although genetic evidence has linked fetal growth with adult metabolism and diabetes risk,^7^ the underlying aetiology remains unclear. Life-long perturbations in causal risk factors, e.g., elevated low-density lipoprotein (LDL) cholesterol, hypertension and type 2 diabetes, might represent pathways by which intrauterine environment affects later cardiometabolic risk.^2,3^ Numerous studies have shown associations between lower birth weight and adverse levels of metabolic risk factors in adulthood; primarily with higher insulin and blood pressure, but also with adverse circulating lipid levels and markers of inflammation.^8-13^ However, meta-analyses have indicated modest magnitudes of association, and controversy prevails regarding the relevance of low birth weight on the circulating lipid profile in adulthood and other cardiometabolic risk markers.^6,8,13,14^ Detailed metabolic profiling in large cohorts can assist in characterizing the molecular aberrations associated with birth weight. This may also clarify the public health importance of metabolic associations with birth weight in comparison to later metabolic effects of adiposity.^15^

The molecular effects of impaired fetal growth involve multiple metabolic pathways which extend beyond routine risk markers.^6^ However, the wider metabolic influences in adulthood have not been studied in large population settings. Metabolomics is a powerful tool to study fine-grained molecular profiles and is therefore an attractive tool to study how birth weight is reflected in the detailed metabolic profile in adulthood. Nuclear magnetic resonance (NMR) metabolomics enables quantitative metabolic profiling of large blood sample collections.^16^ This methodology provides detailed lipoprotein subclass profiling, as well as quantification of fatty acids and small molecules that have recently been linked with the risk for cardiovascular disease and diabetes.^17-21^ These biomarkers could potentially serve as molecular intermediates between impaired intra-uterine growth and cardiometabolic risk. However, only few metabolomics studies on birth weight have been conducted to date. These studies have been limited to a very small number of individuals, have focused on preterm birth at very low birth weight, and primarily assessed associations with metabolites measured from umbilical cord blood.^22-24^ No study has previously used metabolomics to assess the role of birth weight on adult metabolic profiles in large general population settings.

To characterize metabolic signatures of lower birth weight, we used serum NMR metabolomics of 18 288 individuals from five general population cohorts and two twin studies, which together cover individuals from adolescence to the end of working age. We further compared how the metabolic association pattern with birth weight resembles the association pattern of adiposity in adulthood for the same extensive panel of metabolic measures.

## METHODS

### Study populations

The study comprised six Finnish cohorts and one cohort from the United Kingdom (**Table 1**): the children part of the Avon Longitudinal Study of Parents and Children (ALSPAC; n=2874; metabolic profiles measured from blood samples drawn at age 17)^25^; the Northern Finland Birth Cohort (NFBC) 1986 (n=5579; age 16)^26^ and NFBC 1966 (n=5412; age 31)^27^, the Cardiovascular Risk in Young Finns Study (YFS; n=2273; age 24–48)^28^, the FinnTwin studies FT12 (n=767, age 21–25) and FT16 (n=495, age 23–30)^29^, and the Helsinki Birth Cohort Study (HBCS; n=890; age 62–75)^14^. Details of the cohorts related to the present study are described in the **Supplementary Methods**. Birth weight and gestational age were assessed by a midwife, birth medical records or antenatal care. Out of 19 622 eligible individuals with metabolic profiling data, 18 649 had complete data on birth weight, gestational age, and adult body mass index (BMI). Women who were pregnant (n=115) and individuals on lipid-lowering medication (n=246) at the time of metabolic profiling were excluded, leaving 18 288 individuals for the present analysis. All study participants provided informed consent, and study protocols were approved by the local ethical committees.

**Table 1.**
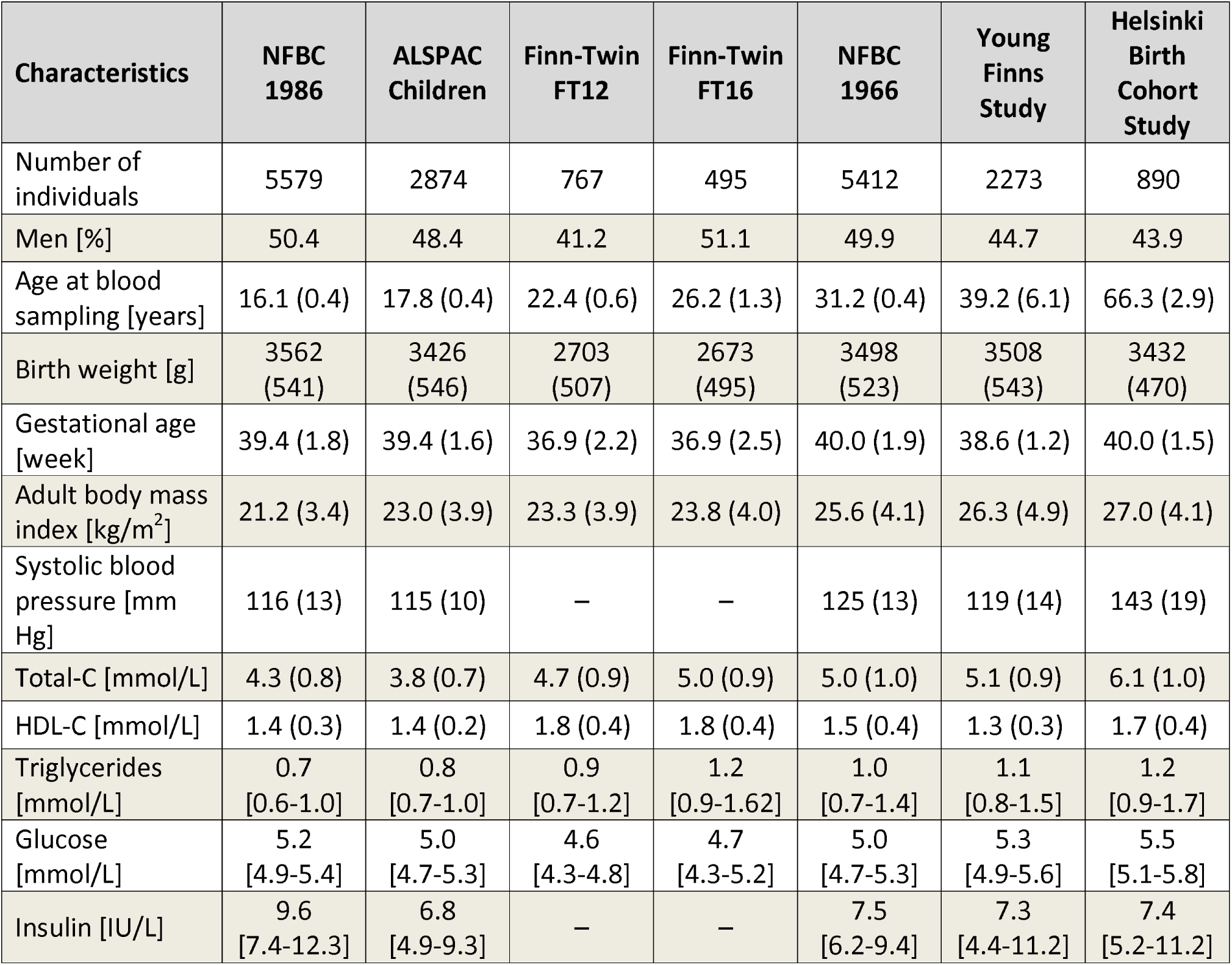
Characteristics of the seven cohorts.

| Characteristics | NFBC<br>1986 | ALSPAC<br>Children | Finn-Twin<br>FT12 | Finn-Twin<br>FT16 | NFBC<br>1966 | Young<br>Finns<br>Study | Helsinki<br>Birth<br>Cohort<br>Study |
| --- | --- | --- | --- | --- | --- | --- | --- |
| Number of individuals | 5579 | 2874 | 767 | 495 | 5412 | 2273 | 890 |
| Men [%] | 50.4 | 48.4 | 41.2 | 51.1 | 49.9 | 44.7 | 43.9 |
| Age at blood sampling [years] | 16.1 (0.4) | 17.8 (0.4) | 22.4 (0.6) | 26.2 (1.3) | 31.2 (0.4) | 39.2 (6.1) | 66.3 (2.9) |
| Birth weight [g] | 3562<br>(541) | 3426<br>(546) | 2703<br>(507) | 2673<br>(495) | 3498<br>(523) | 3508<br>(543) | 3432<br>(470) |
| Gestational age [week] | 39.4 (1.8) | 39.4 (1.6) | 36.9 (2.2) | 36.9 (2.5) | 40.0 (1.9) | 38.6 (1.2) | 40.0 (1.5) |
| Adult body mass index [kg/m<sup>2</sup>] | 21.2 (3.4) | 23.0 (3.9) | 23.3 (3.9) | 23.8 (4.0) | 25.6 (4.1) | 26.3 (4.9) | 27.0 (4.1) |
| Systolic blood pressure [mm Hg] | 116 (13) | 115 (10) | – | – | 125 (13) | 119 (14) | 143 (19) |
| Total-C [mmol/L] | 4.3 (0.8) | 3.8 (0.7) | 4.7 (0.9) | 5.0 (0.9) | 5.0 (1.0) | 5.1 (0.9) | 6.1 (1.0) |
| HDL-C [mmol/L] | 1.4 (0.3) | 1.4 (0.2) | 1.8 (0.4) | 1.8 (0.4) | 1.5 (0.4) | 1.3 (0.3) | 1.7 (0.4) |
| Triglycerides [mmol/L] | 0.7<br>[0.6-1.0] | 0.8<br>[0.7-1.0] | 0.9<br>[0.7-1.2] | 1.2<br>[0.9-1.62] | 1.0<br>[0.7-1.4] | 1.1<br>[0.8-1.5] | 1.2<br>[0.9-1.7] |
| Glucose [mmol/L] | 5.2<br>[4.9-5.4] | 5.0<br>[4.7-5.3] | 4.6<br>[4.3-4.8] | 4.7<br>[4.3-5.2] | 5.0<br>[4.7-5.3] | 5.3<br>[4.9-5.6] | 5.5<br>[5.1-5.8] |
| Insulin [IU/L] | 9.6<br>[7.4-12.3] | 6.8<br>[4.9-9.3] | – | – | 7.5<br>[6.2-9.4] | 7.3<br>[4.4-11.2] | 7.4<br>[5.2-11.2] |

Values are mean (SD) and median [interquartile range] for normally distributed and positively skewed variables, respectively.

### Lipid and metabolite quantification

Fasting blood samples were collected as part of the clinical examinations in adolescence and in adulthood and stored as serum or EDTA plasma at −80°C for subsequent biomarker profiling as detailed in **Supplementary Methods**. Altogether 87 metabolic measures were analysed for the present study. A high-throughput NMR metabolomics platform was used for the quantification of 77 metabolic measures.^16^ This metabolomics platform provides simultaneous quantification of routine lipids, lipid concentrations of 14 lipoprotein subclasses and major subfractions, and further quantifies abundant fatty acids, amino acids, ketone bodies and gluconeogenesis-related metabolites in absolute concentration units (**Table S1**). The metabolic profiling therefore includes both routine risk markers and novel metabolic biomarkers that have not previously been examined in relation to birth weight. The NMR metabolomics platform has been extensively applied for biomarker profiling in epidemiological studies^15,17,18,20,21,30,31^ and details of the experimentation have been described elsewhere.^16,32^ In addition to the NMR metabolomics measures, 10 metabolic markers related to inflammation, liver function and hormone balance, assayed in two or more of the cohorts, were analysed as part of the comprehensive metabolic profile (**Supplementary Methods**).^15^

### Statistical analyses

Metabolic measures with skewed distributions (skewness>2) were normalized by log-transformation prior to analyses. Linear regression models for each metabolite were tested with birth weight as the explanatory variable and the metabolite concentration as an outcome. Associations were adjusted for sex, age at blood sampling, and gestational age. Results were analysed separately for the seven cohorts and combined using fixed effect inverse-variance weighted meta-analysis after verifying the consistency across the seven cohorts. Association magnitudes are quantified in SD-units of metabolite concentration per 1-kg lower birth weight (≈2 SD). Due to the correlated nature of the metabolic measures, >95% of the variation in the 87 measures was explained by at most 34 principal components in each cohort. Multiple testing correction therefore accounted for 34 independent tests using the Bonferroni method, resulting in P<0.0015 denoted statistically significant.

The pattern of metabolic associations with birth weight was compared to the corresponding cross-sectional metabolic associations with BMI, using the same approach as used for birth weight but without adjustment for gestational age. The overall correspondence between the metabolic association patterns of birth weight and adult BMI were summarized using the *R*^2^ and slope of the linear fit.^15,31^

To examine whether there was evidence for curvilinear associations we assessed the shape of the associations using local quadratic regression fitting, with each smoothing function evaluated at 25 points through the range of birth weight. Absolute concentrations of each metabolic measure were first regressed for age and sex, and the resulting residuals were pooled and rescaled to absolute units prior to fitting.^31^

## RESULTS

The study comprised 18 288 adolescents and adults from five general population cohorts and two twin cohorts from Finland and the UK (**Table 1**). Distributions of birth weight for each cohort are illustrated in **Figure S1**. Only 1% of the singleton participants had a birth weight of <2 kg. Birth weight was correlated with adult BMI (*r*=0.09) in a broadly linear manner for most of the cohorts (**Figure S2**).

### Lipoprotein measures

Associations of birth weight with 43 lipoprotein measures are shown in **Figure 1**. In the meta-analysis, birth weight was robustly associated with numerous lipoprotein measures (P<0.0015 for 25 measures). While the associations of routine cholesterol measures were of modest magnitudes, somewhat stronger associations were observed for many of the more detailed lipid measures. Lower birth weight was associated with higher circulating apolipoprotein B (apoB) and total lipid concentrations in the apoB-carrying particles (very-low-density lipoprotein (VLDL), intermediate-density lipoprotein (IDL) and LDL). The strongest associations with lipoprotein subclasses were observed for lipids in medium and small VLDL particles. Associations were weaker for lipids in IDL and LDL particles, albeit with stronger association magnitude for lipids in small LDL. Associations were more heterogeneous for lipids in high-density lipoprotein (HDL) particles: lower birth weight was associated with lower concentrations of circulating lipids in large HDL particles, but with higher concentrations of those in small HDL (i.e., in the same direction as for the apoB-carrying lipoproteins). Birth weight was also robustly associated with the average size of the lipoprotein particles, with increased VLDL size and decreased LDL and HDL size related to lower birth weight. Within a given lipoprotein class, associations with birth weight tended to be stronger for triglycerides than for cholesterol and phospholipid levels.

**Figure 1.**
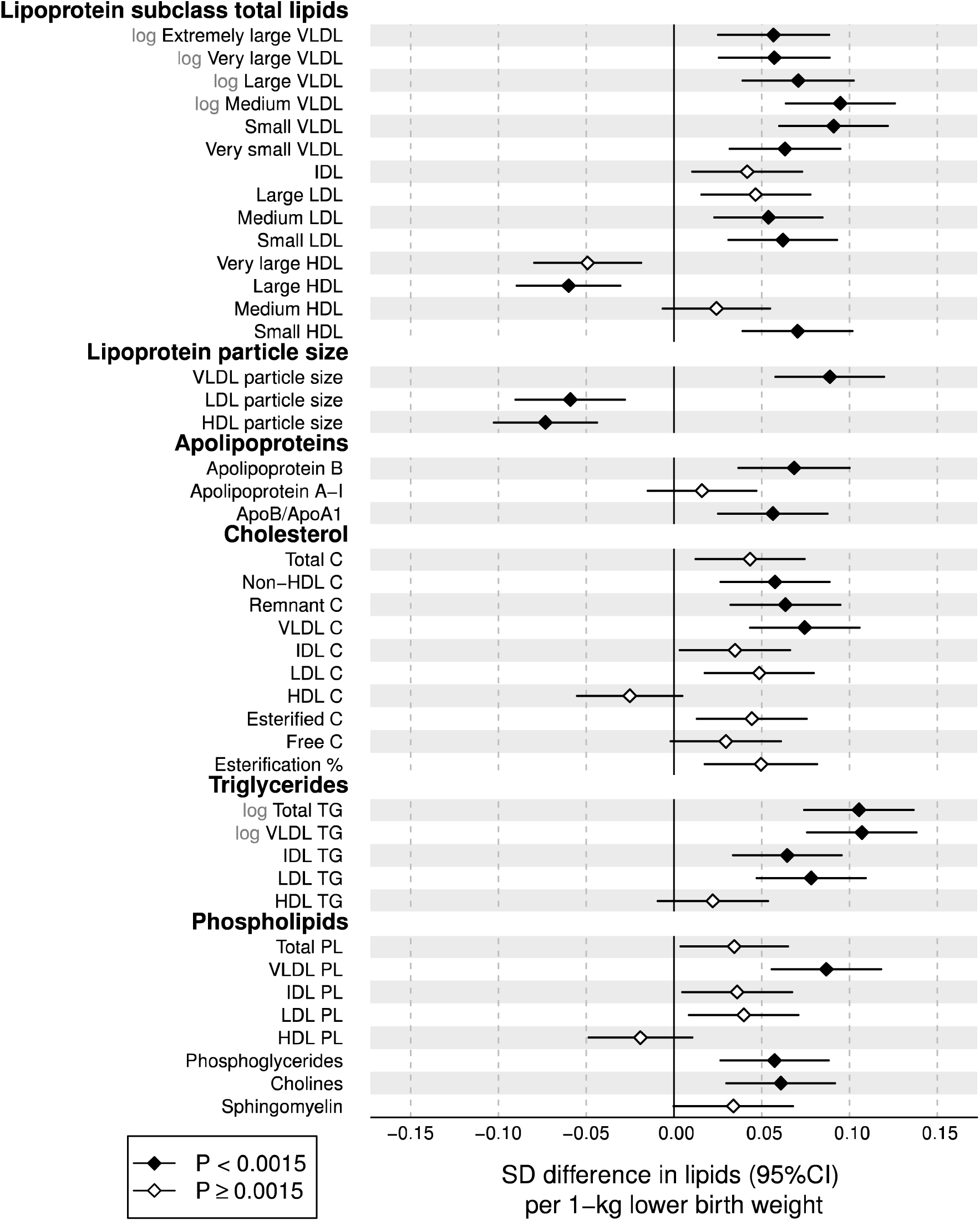
Birth weight associations with adult concentrations of lipoprotein lipids. The associations were adjusted for sex, gestational age, and age at blood sampling, and meta-analysed for 18 288 individuals from 7 cohorts. Association magnitudes are 1-SD lipid concentration per 1-kg lower birth weight. Error bars denote 95% confidence intervals. Filled diamonds indicate P<0.0015. Association magnitudes in absolute concentration units and P-values are listed in **Table S1**. Results for individual cohorts are shown in **Figure S5**.

To enable comparison of birth weight associations across the metabolic measures, all association magnitudes are scaled to SD-units of metabolite concentration per 1-kg lower birth weight. The corresponding associations in absolute units, e.g., mmol/L per kg, are listed in **Table S1**. For instance, total triglyceride concentration was among the measures most strongly associated with birth weight, with each 1-kg lower birth weight being associated with 0.04 mmol/L higher serum triglyceride concentration.

### Fatty acids

Associations of birth weight with 16 fatty acid measures are shown in **Figure 2**. Lower birth weight was robustly associated with higher absolute concentration of all fatty acids assayed except docosahexaenoic acid. The strongest associations were observed for total, saturated and monounsaturated fatty acids (MUFA), which displayed association magnitudes comparable to that of apoB. Somewhat weaker associations were observed for omega-6 and omega-3 fatty acids. For the fatty acid ratios, the proportion of saturated fatty acids and MUFAs tended to be higher among individuals with lower birth weight, whereas the proportion of omega-6 fatty acids were lower.

**Figure 2.**
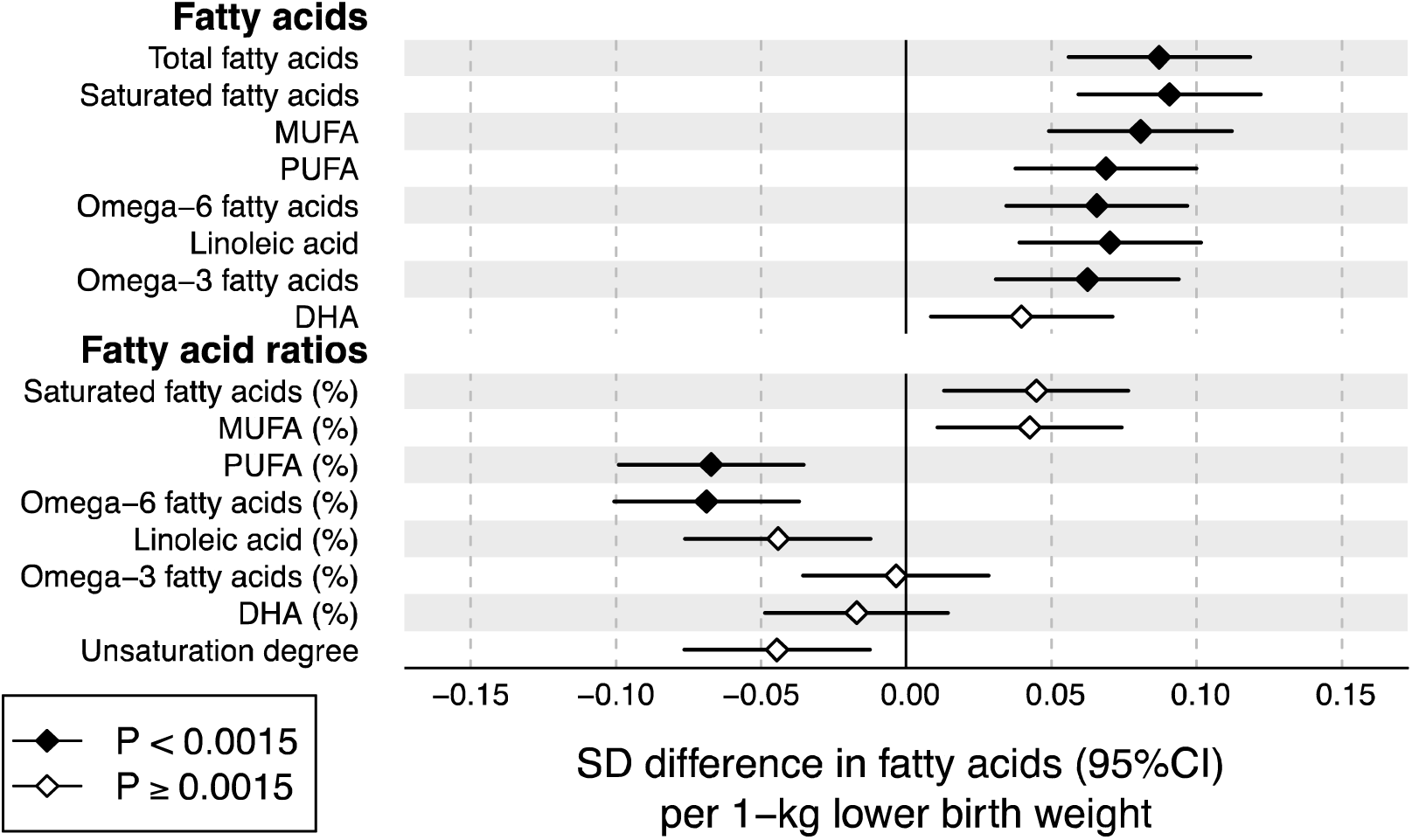
Birth weight associations with adult fatty acid levels. The associations were adjusted for sex, gestational age, and age at blood sampling, and meta-analysed for 18 288 individuals from 7 cohorts. Association magnitudes are in units of 1-SD fatty acid measure per 1-kg lower birth weight. Fatty acid ratios are relative to the total fatty acid concentration. Error bars denote 95% confidence intervals. Filled diamonds indicate P<0.0015. Results for individual cohorts are shown in **Figure S5**. MUFA: mono-unsaturated fatty acids, PUFA: poly-unsaturated fatty acids, DHA: docosahexaenoic acid.

### Non-lipid metabolic measures

Associations of birth weight with 28 non-lipid metabolites and other metabolic measures are shown in **Figure 3**. Lower birth weight was associated with higher concentrations of alanine, branched-chain and aromatic amino acids. These amino acid associations with birth weight were of comparable magnitude to that of apoB. Lower birth weight was not robustly associated with glucose, while other gluconeogenesis-related metabolites displayed stronger associations. Lower birth weight was also robustly associated with higher levels of insulin and certain markers for low-grade inflammation and impaired liver function.

**Figure 3.**
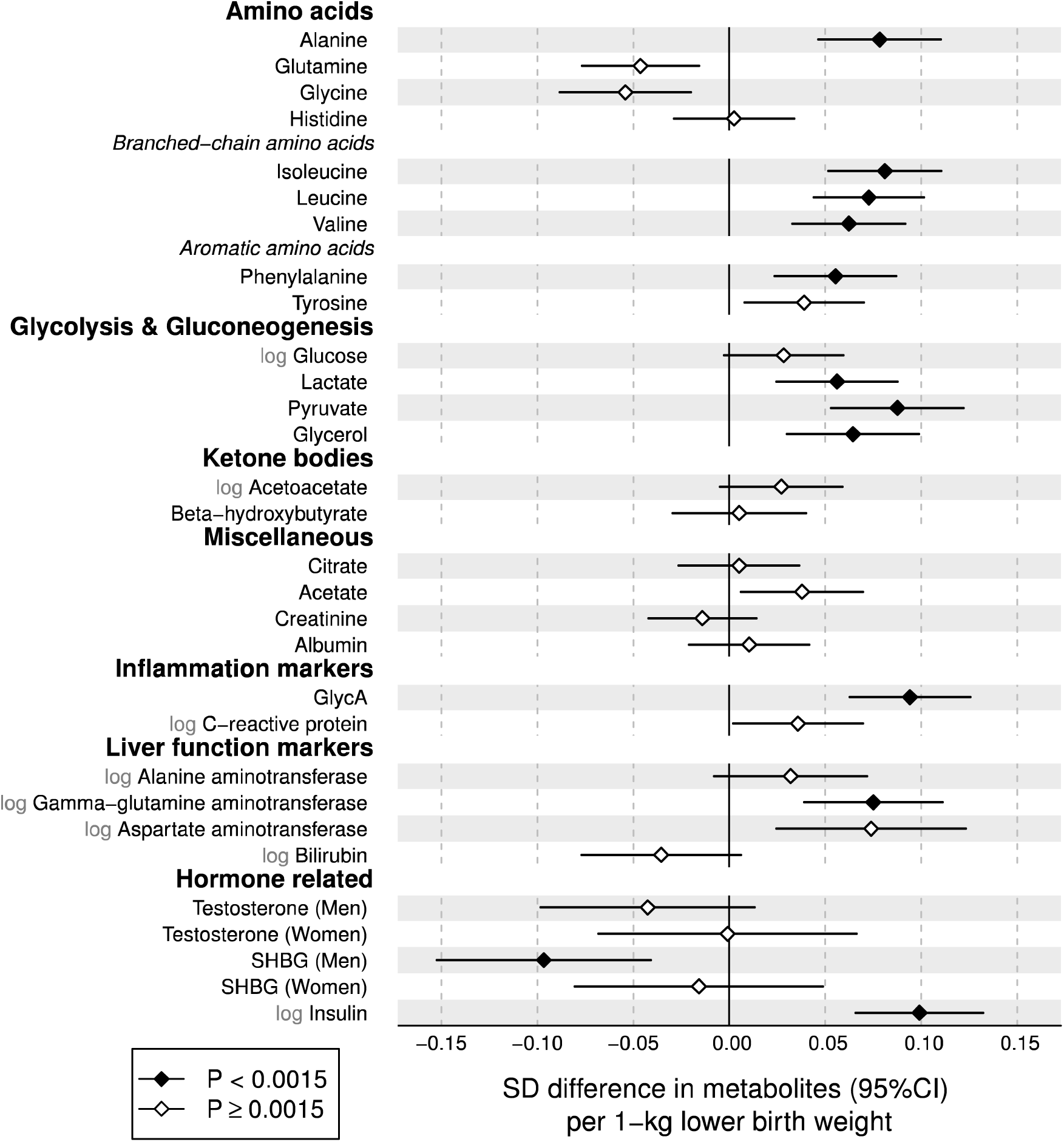
Birth weight associations with adult metabolite and hormonal concentrations. The associations were adjusted for sex, gestational age, and age at blood sampling, and meta-analysed for 18 288 individuals from 7 cohorts. Association magnitudes are in units of 1-SD metabolite concentration per 1-kg lower birth weight. Error bars denote 95% confidence intervals. Filled diamonds indicate P<0.0015. Results for individual cohorts are shown in **Figure S5**. GlycA: Glycoprotein acetyls, SHBG: sex-hormone binding globulin.

### Resemblance between metabolic signatures of birth weight and BMI

The overall pattern of metabolic associations with lower birth weight was reminiscent of the metabolic association pattern with adiposity (**Figure S3**).^9,10,15^ We therefore compared the metabolic associations of birth weight with the corresponding metabolic associations of current BMI (**Figure 4**). The resemblance between the metabolic association patterns was high, as indicated by the goodness-of-fit being *R*^2^=0.77. The slope denotes that 1-kg lower birth weight is associated with similar magnitudes of metabolic aberrations as those linked with 0.92 higher BMI-units (kg/m^2^) in adulthood. A similar resemblance with the association pattern with lower birth weight was observed for higher adult weight (*R*^2^=0.75), indicating that metabolic associations with 1-kg lower birth weight were similar to those linked with 3.1 kg higher adult weight. In contrast, the pattern of metabolic associations with adult height was considerably different (**Figure S4**; *R*^2^=0.17).

**Figure 4.**
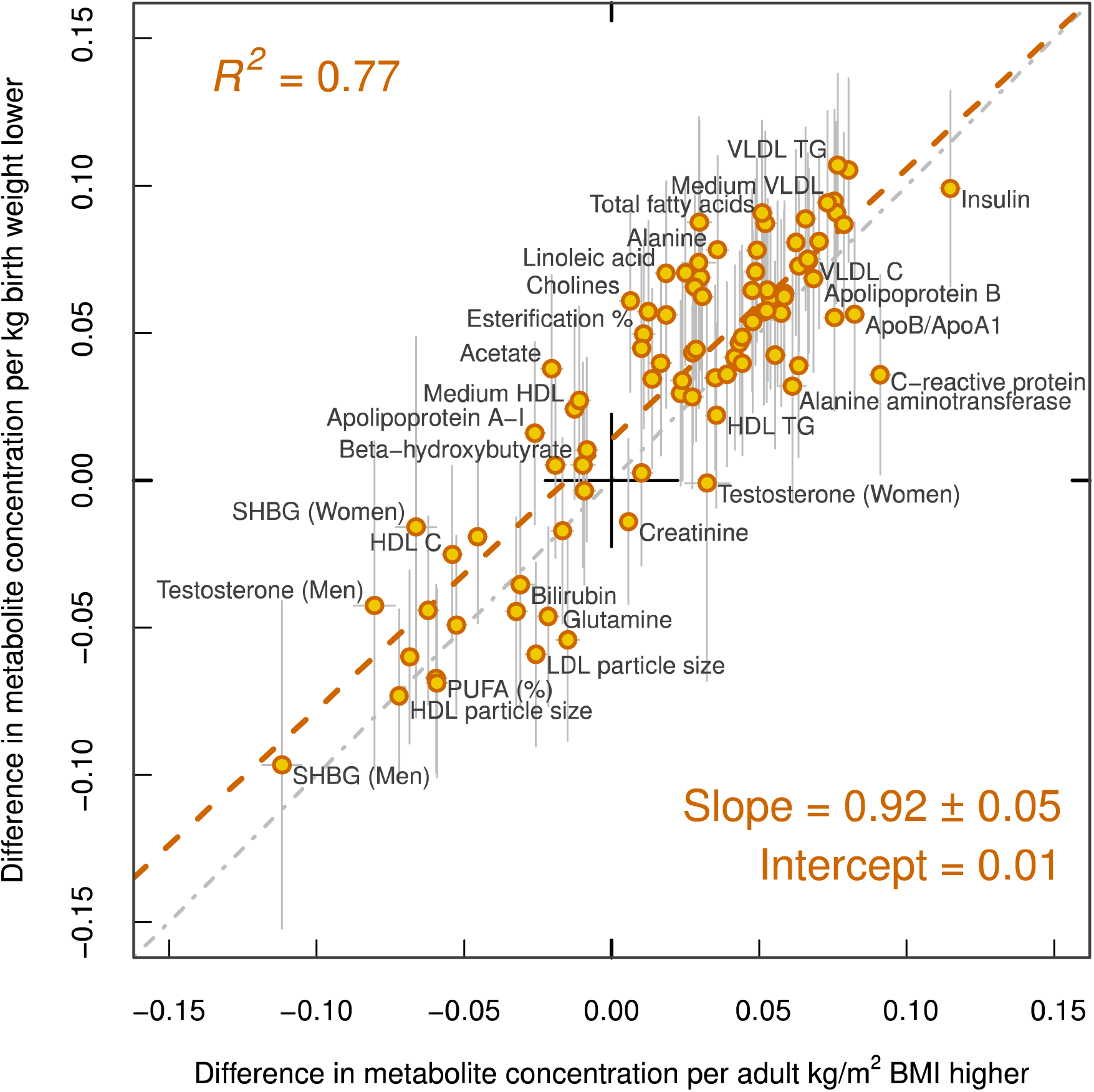
Resemblance between metabolic association patterns related to lower birth weight and higher adulthood BMI. The metabolic associations were assessed for the same 18 288 individuals. The red dashed line denotes the linear fit between metabolic associations with lower birth weight and higher BMI. The slope indicates that 1-kg lower birth weight is on average associated with similar metabolic deviations as those linked with 0.92 kg/m^2^ higher BMI in adulthood.

### Consistency and sensitivity analyses

Despite differences in age at blood sampling, the associations of birth weight with the metabolic measures were consistent across the seven cohorts (**Figure S5**). The results were also generally similar and statistically consistent for men and women (**Figure S6**). The association magnitudes were also similar (on average 5% weaker) if omitting participants within the lowest percentile (birth weight <2.0 kg) and highest percentile (birth weight >4.7 kg). The metabolic associations with birth weight followed a similar pattern if adjusting for adult BMI, but this adjustment increased the magnitude of association by 58% on average (**Figure S7**). The continuous shapes of the metabolic associations with birth weight are illustrated in **Figure S8**. Most associations were approximately linear across the birth weight distribution, justifying the use of linear regression modelling.

## DISCUSSION

In this meta-analysis of 18 288 individuals, lower birth weight was adversely associated with numerous blood-based biomarkers across the comprehensive metabolic profile, including lipoprotein subclass measures, fatty acid composition, amino acids as well as markers of inflammation and liver function. The metabolic associations were all in a direction of higher risk for diabetes and cardiovascular disease, both for established risk factors and emerging biomarkers. The associations were coherent across seven cohorts with a wide age range at metabolic profiling, from adolescence through adulthood, suggesting that these metabolic aberrations are lifelong. Although many of the associations were statistically robust in this large study sample, the small association magnitudes indicate that the influences of birth weight on systemic metabolism are modest at best and likely of limited public health importance.

The wide palette of the metabolic deviations associated with lower birth weight have previously been linked with increased risk for cardiometabolic diseases.^16-21^ Among the lipoprotein measures adversely associated with lower birth weight were both cholesterol-rich LDL particles and triglyceride-rich VLDL particles. Genetic evidence suggests that both these types of apoB-carrying lipoproteins are causally related to ischemic heart disease.^33^ Associations with routine cholesterol measures were modest, in line with prior meta-analyses that have questioned the relevance of associations between birth weight and circulating lipids.^8,34,35^ Although the lipoprotein subclass profiling indicated somewhat stronger and statistically robust associations with several more detailed lipid measures, the associations were nevertheless in line with the overall weak metabolic aberrations.

Increased circulating levels of omega-3 fatty acids have been associated with lower risk for future cardiovascular disease events.^17^ Whilst dietary supplementation of omega-3 fatty acids has been suggested as intervention for people born with impaired fetal growth,^36^ the serum levels of omega-3 have not been robustly linked with birth weight. Here, the association of birth weight was flat for the proportion of omega-3 fatty acids relative to total fatty acids. However, the proportion of omega-6 fatty acids — also inversely associated with the risk for cardiovascular disease^17^ and diabetes^20^— was decreased in relation with lower birth weight. The small magnitudes of the fatty acid perturbations are unlikely to substantially mediate the associations between lower birth weight and increased susceptibility to cardiometabolic diseases.

Recent metabolomics studies have shown that amino acids and many other circulating metabolites are predictive of the risk for diabetes, cardiovascular disease, and all-cause mortality.^17-19,21,37^ For instance, elevated circulating levels of branched-chain and aromatic amino acids have been robustly associated with insulin resistance, hyperglycaemia and diabetes risk in a number of studies.^19,21,37^ Aromatic amino acids have also been shown to be predictors of cardiovascular event risk, even more strongly than LDL cholesterol.^17^ All these amino acids were found to be elevated for lower birth weight, i.e., in the direction consistent with higher metabolic risk. Lower birth weight was also related to higher concentrations of markers for impaired liver function, insulin resistance and chronic inflammation. These metabolic markers are also predictive for the risk of a broad span of chronic diseases and all-cause mortality.^18,38,39^ Importantly, while the individual biomarker associations with birth weight are all modest, the combined metabolic aberrations may in concert potentially contribute to mediate the relation between birth weight and cardiometabolic disease risk.

Elevated BMI has recently been shown to have a causal metabolic signature across biomarkers from multiple pathways.^15^ This metabolic signature of adiposity is highly reminiscent of the pattern of metabolic associations here linked with lower birth weight. The similarity in the detailed association patterns illustrates how comprehensive metabolic profiling may help to pinpoint molecular connections between the metabolic effects of different risk factors. The resemblance between the metabolic signatures allows to summarize the metabolic aberrations linked with birth weight in relation to the effects of adiposity: 1-kg lower birth weight in the general population is associated with similar metabolic deviations as those caused by 0.92 kg/m^2^ higher BMI in adulthood. The consistency with the metabolic association pattern for birth weight was driven by adult weight rather than height, suggesting that the underlying mechanisms are not primarily related to growth. A 1-kg difference in birth weight is substantial, corresponding to almost 2 SDs. A similar variance in adulthood BMI (i.e., a 2 SD greater adult BMI) is associated with 8-fold stronger metabolic deviations, thus giving some perspective to the public health relevance of the diverse metabolic perturbations associated with lower birth weight.

The clear correspondence between the metabolic signatures of lower birth weight and elevated BMI led us to hypothesize that impaired fetal growth and adiposity may have shared molecular pathways underlying the fine-grained metabolic aberrations. As a corollary of this hypothesis, we anticipate that the causal effects of BMI on also other molecular markers^40^ and physiological factors can be used to predict the anticipated association of birth weight with these same outcomes. For instance, genetic evidence indicates that the lifelong causal effect of BMI on systolic blood pressure is 0.9 mmHg per higher BMI-unit;^15,41^ based on the above hypothesis we extrapolate that the association of birth weight with systolic blood pressure would be 0.9×0.92 ≈ 0.8 mmHg per 1-kg lower birth weight. This is broadly consistent with the association observed in meta-analysis.^9,10^ Since our results indicate that the metabolic perturbations associated with birth weight are present across the lifecourse, it is important that such extrapolation of birth weight associations with other risk markers is based on lifelong effects of BMI, e.g., from genetic estimates. Based in this principle, it is also possible to estimate the association of birth weight with cardiometabolic disease risk by comparison to the risk effects caused by BMI. Accordingly, 0.92 BMI-units is causally associated with ≈10% higher risk for ischemic heart disease,^42,43^ which is consistent with meta-analysis results on the risk magnitude for ischemic heart disease per 1-kg lower birth weight.^2^ Similarly, 0.92 BMI-units is causally associated with 24–33% higher risk for type 2 diabetes^41,44^, which again is consistent with meta-analysis estimates of the diabetes risk per 1-kg lower birth weight.^3^ These results seem to suggest that the comprehensive metabolic effects corresponding to as little as 0.92 BMI-units, affecting over the lifecourse, may be sufficient to explain the association of birth weight with cardiometabolic disease risk in adulthood.

It is of note that although birth weight was ≈700 grams lower in the twin cohorts than in singletons, the metabolite concentrations and the pattern of metabolic associations with birth weight were similar to the other cohorts. This is consistent with no difference in diabetes prevalence or overall mortality among twin individuals compared to singletons ^45,46^, supporting our conclusion that lower birth weight per se has a minor long-term impact on the systemic metabolic risk profile. The metabolic associations with birth weight were stronger if adjusting for adult BMI, suggesting that birth weight in relation to adult adiposity is more relevant for metabolic risk than birth size alone.^47^ Further studies on the comprehensive metabolic effects of growth in infancy may clarify the role of compensatory growth and other proposed interactions with birth weight on the metabolic profile in adulthood.^14^

Strengths of this study include the large sample size, comprising seven cohorts with quantitative metabolomics data. Birth weight was obtained from birth medical records for 88% of the study population, which minimizes bias from self-reporting, and data on gestational age allowed accounting for lower birth weight caused by prematurity. The broadly coherent results across cohorts of a wide age range provided a view to the life-course effects of birth weight. However, the general population nature of the cohorts analysed prevents us from making conclusions regarding the specific metabolic effects of infants born preterm, or those with severe fetal growth restriction.^24^ Birth weight has limitations as a marker of impaired fetal growth, however it is the surrogate most widely reported in large population cohorts, and it has a high correlation with other markers of size at birth.^35^ We acknowledge that we cannot assume that impaired fetal growth is causal for the adulthood metabolic aberrations observed. For example, genetic analyses support a causal role for maternal smoking in pregnancy and higher blood pressure resulting on lower infant birth weight, which could potentially contribute to explain the weak inverse associations with metabolic risk markers observed here.^48,49^ Given the predominantly young age of the study participants we were not able to test whether the metabolic aberrations related to birth weight could mediate the relationship to cardiometabolic disease outcomes. Finally, it is important to recognize that information on antenatal nutrition and other factors that might underlie the relationships of birth weight with metabolic outcomes might highlight associations of a larger magnitude or potentially greater public health importance than suggested by our results.

In conclusion, comprehensive metabolic profiling of large cohorts identified associations between lower birth weight and adverse circulating levels of a wide panel of circulating cardiometabolic risk markers in adulthood. The overall metabolic signature of lower birth weight closely resembled the metabolic effects of higher adiposity, suggesting that shared molecular pathways may underpin the perturbed metabolic profile. Nevertheless, the aberrations were of modest magnitude, with similar metabolic perturbations related to 1-kg lower birth weight as those caused by lifelong effects of ≈3 kg higher body weight in adulthood. These results indicate that birth weight is only a weak indicator of metabolic risk in adulthood.

## Supplementary Data

**Supplementary Methods: Study populations**.

**Table S1. Mean (SD) metabolic concentrations, and associations with birth weight in absolute concentration units**.

**Figure S1. Birth weight distribution in each cohort**.

**Figure S2. Adulthood body mass index as a function of birth weight in each cohort**.

**Figure S3. Metabolic associations with adulthood body mass index**.

**Figure S4. Metabolic associations with adulthood height**.

**Figure S5. Metabolic associations with birth weight in each cohort**.

**Figure S6. Metabolic associations with birth weight separately for men and women**.

**Figure S7. Metabolic associations with birth weight when omitting adjustment for gestational age and BMI**.

**Figure S8. Curvilinear shapes of metabolic associations with birth weight**.

